# Circuit motifs and graph properties of connectome development in *C. elegans*

**DOI:** 10.1101/2021.07.11.451911

**Authors:** Jordan K. Matelsky, Raphael Norman-Tenazas, Felicia Davenport, Elizabeth P. Reilly, William Gray-Roncal

**Author notes:** Correspondence: {, }.

## Abstract

Network science is a powerful tool that can be used to better explore the complex structure of brain networks. Leveraging graph and motif analysis tools, we interrogate *C. elegans* connectomes across multiple developmental time points and compare the resulting graph characteristics and substructures over time. We show the evolution of the networks and highlight stable invariants and patterns as well as those that grow or decay unexpectedly, providing a substrate for additional analysis.

## Introduction

As new synapse-resolution connectomics datasets are generated at increasing spatial volume extents, the neuroscience community is now faced with the exciting challenge of interpreting and exploring datasets that are much too large and numerous to manually proofread or inspect at the neuron-level. Network analysis tools are one powerful strategy because they enable us to interpret the brain as a graph, where neurons are represented by nodes, and synapses are represented by directed edges. This simplified representation of connectivity provides one approach for summarizing important graph circuits and network properties. We leverage existing technologies and software, such as graph analysis toolkits and graph databases, in order to study the brain for advances in computation, health, and disease (1–3).

The *C. elegans* nematode serves as a well-studied, small nervous system thought to be highly conserved across isogenic individuals. Recent efforts (4) have mapped the complete *C. elegans* “wiring diagram,” or *connectome*, of individuals at different ages, which provides us with a rich and useful set of graphs for comparison. This multi-individual dataset is a powerful tool to understand the development, modularity, and structure of a complete connectome throughout maturation and across a population, foreshadowing future studies in larger organisms such as *Drosophila*, mice, and humans. We leverage network science approaches in order to better characterize the types of modifications that the *C. elegans* brain undergoes throughout development.

## Results

We identified and leveraged several graph analysis tools commonly used by the community to evaluate and explore connectome networks. We performed both computationally-inexpensive graph summary analyses, as well as more computationally-expensive subgraph structure analyses in order to better understand local connectome graph topology.

We share a selection of reproducible and noteworthy findings on seminal nematode connectomes from recent years (4, 5).

### Undirected motif atlas scans of the growing *C. elegans* connectome reveal a changing local structure

We replicated the motif atlas-scan methods previously applied to partial mouse and invertebrate connectomes (7). This atlas-scan procedure involves searching for and counting all undirected subgraphs up to a certain size (here, subgraphs with six vertices or fewer). There are 210 such graphs. **Fig. 1** shows the absolute motif counts of each motif for each of the eight datasets in (4).

**Fig. 1.**
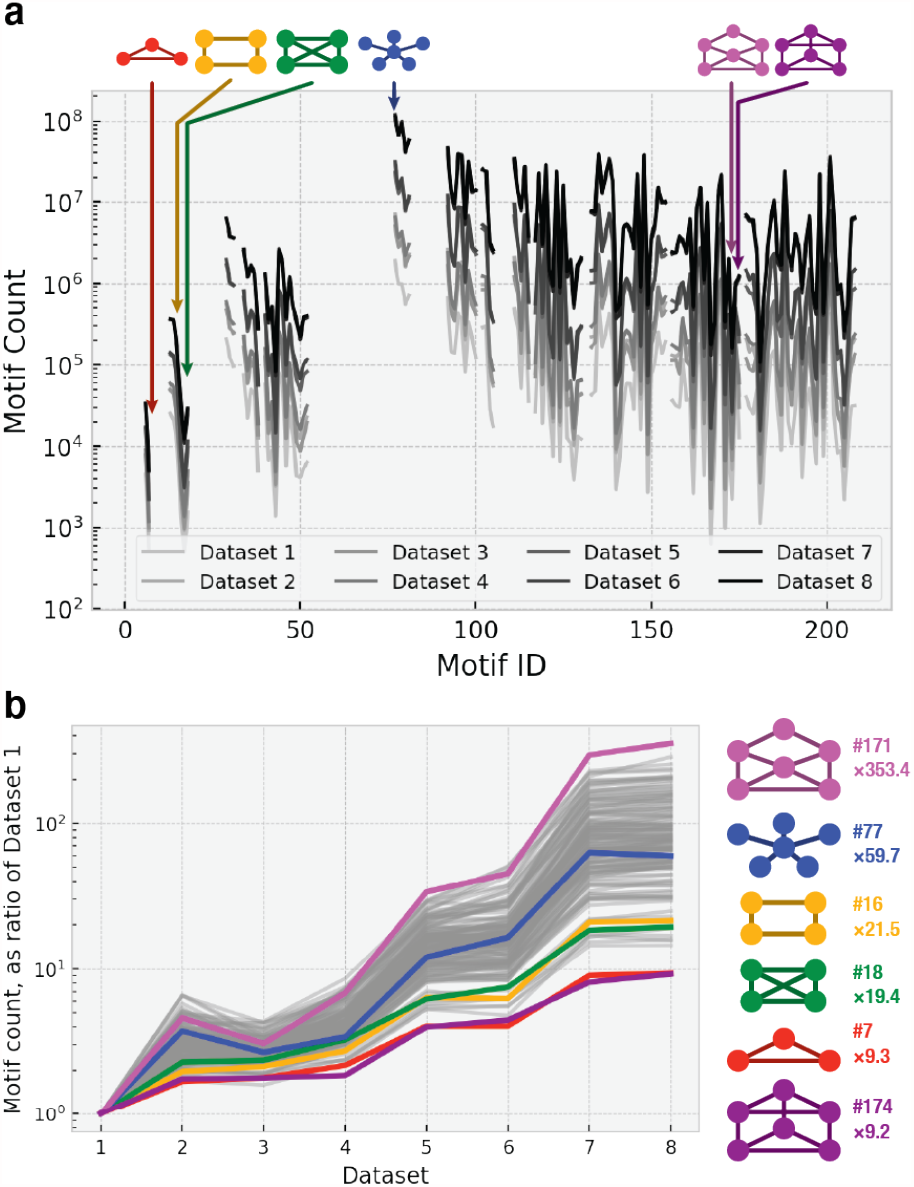
Connectome motif growth over the lifespan of *C. elegans*. **a**. Undirected motif prevalence, showing all undirected graphs with six or fewer neurons. Six graphs of interest are identified along the *x* axis. **b**. Motif growth throughout development. Motif counts are shown as a ratio of counts in Dataset 1: That is, a *y*-value of 2 indicates that there are twice as many of that motif as there were in Dataset 1. All undirected motifs from panel **a** are shown in grey. The six motifs of interest are rendered in color. The most rapidly growing motif (magenta) occurs 353 times more frequently in Dataset 8 as in Dataset 1. The most slowly growing motifs are triangles (red) and the triangular prism net (purple), at 9.24× and 9.33× prevalence in Dataset 8. The motifs shown are identified in the *Atlas of Graphs* (6) as IDs 171, 77, 16, 18, 7, and 174 (from top to bottom).

We then related the motif prevalence at each timestep to the prevalence at the first timestamp, Dataset 1 (**Fig. 1b**). This gives us a measure of the growth of motifs over developmental time. We call particular attention to the slowest-and fastest-growing motifs, as well as other motifs of interest. Using IDs from the *Atlas of Graphs* (6): Motifs #174 and #7, the triangular prism net and the triangle graph, are the slowest-growing motifs, with only a nine-fold increase in motif prevalence. Motif #171 is the fastest-growing motif, with more than a 350-fold increase in motif prevalence between datasets 1 and 8. This is particularly notable in light of the substantial structural similarities between this motif and the slowest-growing #174. Motif #77, the motif with the highest absolute count at all eight timesteps, is of average growth-rate, suggesting that absolute motif count and motif growth rate are *not* closely related measures.

### Neuron degree increases during development

The average and maximum degree of a neuron in the *C. elegans* connectome tend to increase with animal age (**Fig. 3**). This is in contrast with the pruning process known to be common to vertebrate connectomes, in which high-degree neurons eliminate unnecessary synapses during early development. Because animal age and certain connectome statistics — such as maximum neuron degree or edge count — appear to be highly correlated, it may be possible to estimate such properties from an animal age alone. All neurons’ degrees increase with age (**Fig. 2**). The only cell with a final degree smaller than its respective degree in Dataset 1 is *GLRVL*, a muscle-derived glial cell (9) — a result that may be attributable to biological noise.

**Fig. 2.**
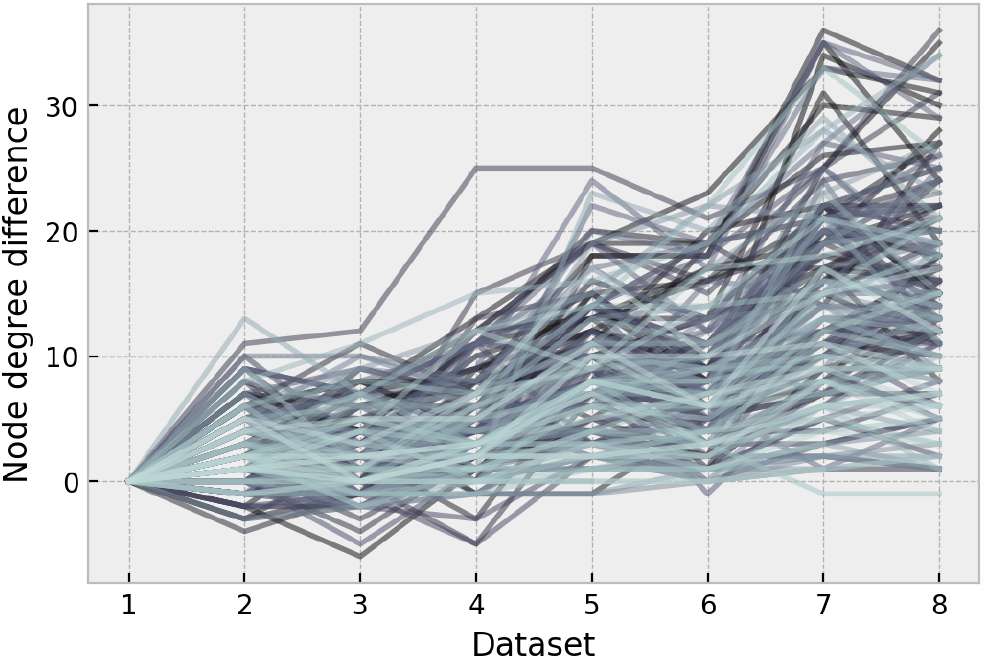
Change in node degree. All changes in node degree are plotted as a function of animal age. The starting degree is defined as the node degree in Dataset 1 (4). The *y*-value is defined as the change in node degree since Dataset 1.

**Fig. 3.**
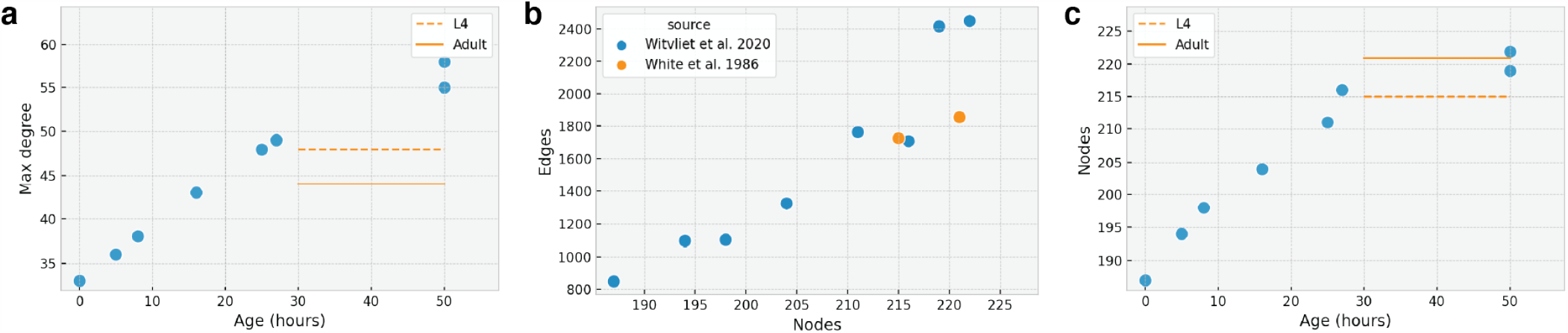
**a**. The average node degree of a connectome increases with age, and is variable across adults. Orange lines denote original datasets for which a precise age was not provided (5). **b**. Node and edge counts in all individuals. **c**. Growth of new neurons over time.

### Graph properties of a growing brain. (Table 1)

We confirm the assertion in Ref. (4) that the adult graphs from Ref. (5) do not have the same properties as the recent adult graphs in Ref. (4). Furthermore, we also confirm that the two new adult datasets (4) have distinct graphs from each other, with distinct graph properties.

**Table 1.** Graph Properties. A table of graph summary statistics for ten connectomes of interest. There are no orphan nodes and no lone pairs. All graphs were weakly connected. Max SCC refers to the number of nodes in the largest strongly connected component.

| Dataset | Nodes | Edges | Loops | Leaves | Max Degree | Mean Degree | Max SCC | Larval Stage | Age (Hours) | Source |
| --- | --- | --- | --- | --- | --- | --- | --- | --- | --- | --- |
| Witvliet 2020 Dataset 1 | 187 | 849 | 2 | 10 | 33 | 9.08 | 121 | L1 | 0 | (4) |
| Witvliet 2020 Dataset 2 | 194 | 1095 | 2 | 8 | 36 | 11.28 | 135 | L1 | 5 | (4) |
| Witvliet 2020 Dataset 3 | 198 | 1101 | 2 | 17 | 38 | 11.12 | 138 | L1 | 8 | (4) |
| Witvliet 2020 Dataset 4 | 204 | 1324 | 3 | 11 | 43 | 12.98 | 149 | L1 | 16 | (4) |
| Witvliet 2020 Dataset 5 | 211 | 1763 | 6 | 5 | 48 | 16.71 | 161 | L2 | 25 | (4) |
| Witvliet 2020 Dataset 6 | 216 | 1707 | 2 | 5 | 49 | 15.80 | 160 | L3 | 27 | (4) |
| Witvliet 2020 Dataset 7 | 222 | 2450 | 16 | 1 | 58 | 22.07 | 191 | Adult | 50 | (4) |
| Witvliet 2020 Dataset 8 | 219 | 2416 | 8 | 2 | 55 | 22.06 | 174 | Adult | 50 | (4) |
| White 1986 JSH | 215 | 1725 | 2 | 2 | 48 | 16.04 | 168 | L4 | – | (5) |
| White 1986 N2U | 221 | 1855 | 2 | 7 | 44 | 16.78 | 170 | Adult | – | (5) |

### The *C. elegans* connectome is dominated by a single strongly connected component

The *C. elegans* connectome is composed of a dense network with inputs predominantly from sensory neurons and outputs predominantly to motor neurons. The (directed) density of the connectomes in Ref. (4) doubles from 0.245 (Dataset 1) to 0.489 and 0.502 in Datasets 7 and 8. The largest strongly connected component contains nearly all neurons, regardless of animal age.

### The adult connectome has significant variability in loop counts

Previous work (10, 11) has suggested that loops and cycles are important for computation in the growing nematode connectome. Here, we use our reproducible graph techniques to quantify this property during development, and show that the total number of loops tends to grow as the animal matures.

### Degree distribution is nonrandom

We compared degree distributions between connectomes, as well as between connectomes and an Erdős–Rényi random graph, following the strategy employed in a recent analysis (2). We show that none of the connectome variants have a degree distribution similar to that of a random graph (**Fig. 4**) by using a Kolmogorov-Smirnov test (*p* ≈ 0).

**Fig. 4.**
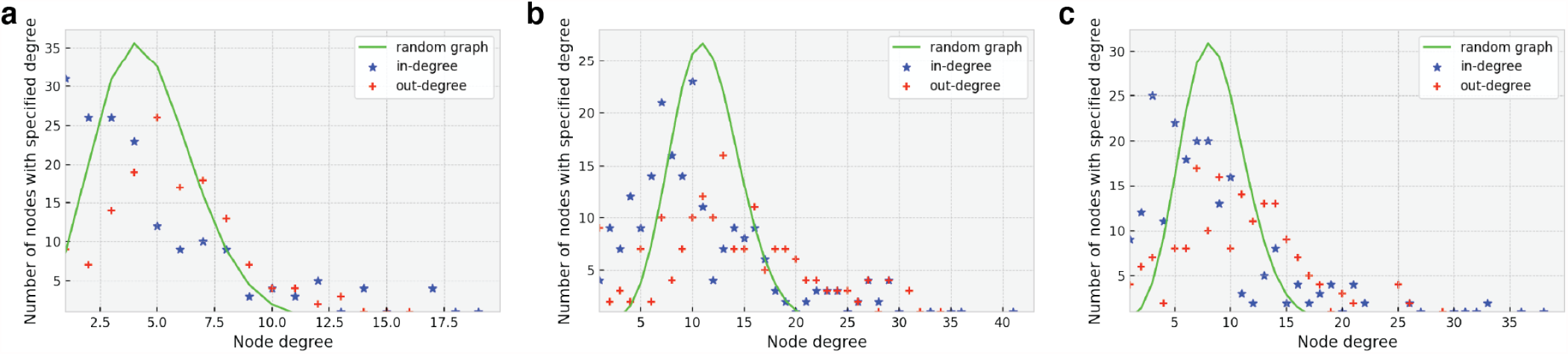
In and out degree frequency distribution of selected connectomes. Random graph is created with the Erdős–Rényi model (8), where the probability of connection is equal to the compared graph’s density. **a**. L1 larvae (4). **b**. Adult connectome (4). **c**. Adult connectome (5). In the adult connectomes, we see that there are a significant number of low-degree and high-degree nodes than random.

## Methods

Taking advantage of previously-developed graph analysis frameworks (7), we computed motif searches (i.e., subgraph isomorphism counts), highlighting the prevalence of interesting structures over development. To understand the macro-structure of the *C. elegans* connectome networks during growth, we employed computationally inexpensive summary statistics, which are increasingly common in the network neuroscience community (12). Despite their apparent simplicity, these measures enable us to better understand the structure and nature of a brain during development.

We computed connectome graph analyses and summary statistics, such as those in **Table 1**, using a scalable network science toolkit developed to operate efficiently on thousand- to billion-edge connectomes. For source code, see *Supplemental Materials*. All analyses were performed on consumer laptop hardware and can scale easily to compute and cluster resources for larger datasets.

## Discussion

We can rapidly distill and analysis key network properties from complex datasets across an organisms development, using scalable, reproducible tools. Future work will include the analysis of sub-circuits within the connectome to see how their structure and function develop over time. An analysis using the volumetric image and segmentation data (4) could further elaborate on differences between adult connectomes and provide more insight into where and how the connectome differs between isogenic individuals.

It is interesting that different undirected subgraphs undergo different rates of growth in the maturing connectome. What biological mechanisms cause such dramatic differences in motif growth-rate? This points to a “preference” for certain structural properties in immature connectomes, and different local structural property preferences in a mature connectome — compatible with parallel observations on topological changes to a connectome during maturation (11).

We hope this approach may serve as a model for the study of larger and less well-understood connectomes, such as emergent fruit-fly, rodent, and human connectomes (13–16).

## Supplemental Materials

Motif-search tools are available at github.com/ aplbrain/dotmotif. Graph algorithm scaling tools are available at github.com/aplbrain/grand. Other software and data will be available upon publication.

## ACKNOWLEDGEMENTS

Research reported in this publication was supported by the National Institute of Mental Health of the National Institutes of Health under Award Numbers R24MH114799 and R24MH114785. The content is solely the responsibility of the authors and does not necessarily represent the official views of the National Institutes of Health. This work was supported by JHU/APL Internal Research Funding.

